# Effects of Social Housing on Electrically Stimulated Dopamine Release in the Nucleus Accumbens Core and Shell in Female and Male Rats

**DOI:** 10.1101/2025.02.25.640103

**Authors:** Ivette L. Gonzalez, Jill B. Becker

## Abstract

Dopamine (DA) is a neurotransmitter that is important in the reward system and increased DA release is associated with rewarding properties of drugs. Highly addictive stimulants like methamphetamine (METH) increase DA release and block reuptake, causing the DA to stay in the synapse longer, enhancing its effects. Because the misuse of METH is increasing in the United States, it is important to investigate ways to protect against this highly addictive stimulant. Recent studies have shown that social support can be a protective factor against METH self-administration in females, but not males. Other studies using microdialysis have shown that socially housed females have lower DA release in the nucleus accumbens (NAc) compared to single housed females after treatment with METH. Additionally, researchers have shown that there are sex differences in stimulated DA release. The present study investigates whether social housing affects stimulated DA release after METH treatment. DA release in the NAc core and shell of socially housed and individually housed rats was measured using fast scan cyclic voltammetry (FSCV) with a chronic 16-channel carbon fiber electrode in the NAc. A stimulating electrode was aimed at the ventral tegmental area (VTA) to induce DA release in the NAc. The results showed that social housing enhances electrically stimulated DA release in males and that there was greater DA release in NAc core than shell in single males, but no difference in socially housed males. In females, social housing also enhanced ES DA release. In single females there was greater ES DA release in shell than in core. Additionally, in single housed females there was greater ES DA release over time, while the socially housed females had high ES DA release that remained stable over time. These results suggest that social housing protects females from sensitization, making single females more vulnerable to the addictive properties of METH.

## Introduction

In recent years, stimulant use has been on the rise. In Massachusetts, for example, stimulant use from 2014 to 2021 ranged from 4% to 7% of the population [1]. More specifically, methamphetamine (METH) use has had a significant rise in recent years [2]. METH acts in the brain by increasing the release of dopamine (DA) and blocking DA reuptake, allowing the released DA to stay longer in the synaptic cleft [3]. METH also has neurotoxic effects caused by neuronal apoptosis, astrocytosis, and microgliosis as well as hyperthermia [3], so its increased use is a public health concern.

In laboratory rats, social housing attenuates cocaine-taking behavior in females but not in males [4]. Female rats who were single housed have higher METH usage compared to females who are socially housed with a naïve partner [5]. Using in vivo microdialysis, individually housed females show a significantly greater increase in DA in the nucleus accumbens (NAc) after METH, compared to socially housed females, [5].

There are also sex differences in stimulated DA release in the NAc and dorsal striatum [6–10]. In previous studies we have successfully used microdialysis to repeatedly measure stimulated DA increases in the NAc [11]. There are significant challenges, however, with using microdialysis repeatedly. In fact, various techniques to directly measure DA release in vivo have struggled with spatial and/or temporal resolution [12].

Fast-scan cyclic voltammetry (FSCV) uses a carbon fiber electrode to measure oxidizable molecules, such as DA with high temporal and spatial resolution [13–16]. FSCV has been used in vitro with brain slices as well as in vivo in anesthetized or freely-moving animals [13–15,17]. Recordings from chronically implanted FSCV electrodes have been used to allow for studies with a more longitudinal design but introduces different challenges such as reduced sensitivity due to immune response [18]. While there have been many advancements in carbon fiber electrode technology for use in freely moving animals, there are still limitations.

In this project we use a novel chronically implanted FSCV electrode to detect DA release in freely moving animals. This 16-channel high density carbon fiber array was designed and manufactured at the University of Michigan [19]. These probes maintain reliable high yield recordings for at least two months [20]. With these electrodes we can simultaneously record from anatomically distinct subregions of the NAc, the core (NAcC) and shell (NAcS). We have measured electrically stimulated (ES) DA release and DA reuptake in the NAcC and NAcS as well as in the dorsal striatum in freely moving animals for weeks [21]. The location of the probe can be determined post-mortem using slice-in-place technology [20].

In previous studies using microdialysis in the NAc, our probes were sampling from both NAcC and NAcS. With the probes used here, we investigate the effect of pair-housing on ES DA release and can now specifically determine the effects in the NAcC and NAcS in male and female rats. This allows us to determine whether we find effects of social housing and/or sex differences when NAcC and NAcS are examined specifically.

## Methods

10 male and 12 female Sprague-Dawley rats approximately 40-45 days old were obtained from Charles River Breeding Laboratory (Portage, MI). Animals were maintained on a 14:10 light/dark cycle in a temperature-controlled climate of 72 ± 2 °F, in ventilated laboratory cages. Rats had ad libitum access to water and phytoestrogen-free rat chow (2017 Teklad Global, 14% protein rodent maintenance diet, Harlan rat chow; Harlan Teklad). All procedures were performed according to protocols approved by the University of Michigan Institutional Animal Care and Use Committee.

Of the total animals, 6 females and 5 males were individually housed. 6 of the females and 5 of males were housed in pairs (social housing) for three weeks before surgery. One of the paired rats went through surgery and was individually housed for 6 days for post-operative care. On day 6, pairs were reunited and remained paired until the end of the experiment. This design was implemented equally for both males and females. Rats were about 75 days old on their first day of FSCV testing (Figure 1).

**Figure 1.**
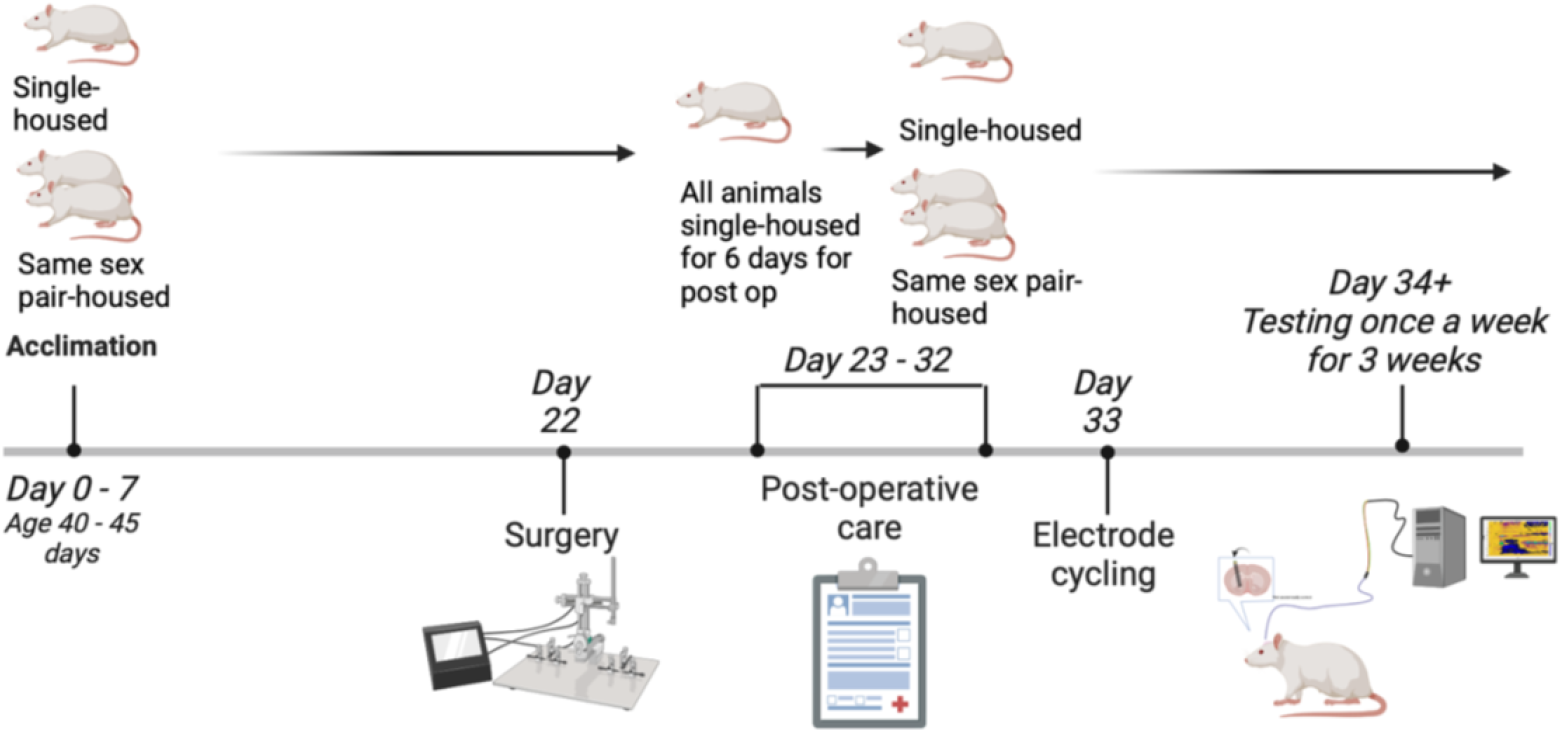
Experimental procedure timeline. (Created with BioRender.com)

### FSCV Surgery

Data was acquired and electrodes calibrated as described previously[21]. A complete guide to the system’s hardware and software specifications can be found at https://chestekresearch.engin.umich.edu/carbon-fibers/. The 16-channel array had a 50μm laser-ablated tip sites prepared as described previously [20]. Devices were sterilized with ethylene oxide. Animals were anesthetized with a combination of ketamine (60 mg/kg, i.p.) and dexmedetomidine hydrochloride (0.3 mg/kg i.p.). A reference electrode cannula (BASi, MD-2251), was prepared as described previously [21] and lowered into the brain (AP -3.0 mm, ML ±3.5 mm, DV (relative to the skull) -1.2 mm). A bipolar stimulating electrode was lowered from the top of the skull into the ventral tegmental area (VTA) (AP -5.15-5.2 mm, ML ±0.75-0.8 mm, DV (relative to the skull) -8.2-8.4 mm). The working electrode tips were brought to the brain surface, the stimulating electrode was lowered 0.2 mm /10 sec while the HDCF array was slowly lowered 0.1 mm every 10 sec with the system on. The final location in the NAc was AP +1.35-1.4 mm, ML ±2.9-3.2 mm, DV (relative to the cortical surface) -6.8-7.2 mm, at an angle of 16.7 degrees. Electrical pulses to the VTA were used to stimulate DA release to insure the working electrode was detecting DA once in place. The working and stimulating electrodes were anchored in place with dental acrylic and the entire headstage was covered with protective dust caps (Omnetics, A79041-001 and Plastics One, Inc., 303DCMN), the reference electrode cannula was occluded with the provided sterile stylet. At the conclusion of the surgery, rats were given atipamezole hydrochloride (0.5 mg/kg i.p.) and dexamethasone (200μg/kg s.c.). The rats then went through standard post-operative care.

### FSCV Testing

At 11 days post-surgery the FSCV electrodes were conditioned as described previously [21], referred to as a ‘cycling session’ which stabilizes the working electrode signal (Figure 2). On testing days, a cycling session occurs before any recordings were taken, and then a 10-minute ES DA signal is collected with the applied waveform repeatedly cycled at a rate of 10Hz. Following the ES DA collection, three 30-second stimulation events recordings were taken, with 5 min between each recording. Five seconds into each recording, a ES pulse train was applied to the stimulating electrode using a stimulus isolator controlled by custom LabVIEW software. Three stimulation parameters were used: pulse trains of 30Hz 15 total pulses (p), 60Hz 30p, or 60Hz 60p. During the initial stimulation test session, the peak amplitude of the current applied was optimized for each rat, ranging from 120-200 μA, and held constant for all pulse trains (Example stimulation parameter: 30Hz 15p, 60Hz 30/60p 150 μA). After the initial stimulations, an intraperitoneal (i.p.) injection of 0.5 mg/kg METH or equivalent volume of saline injection was given. After a 5 min waiting period, DA is recorded for 5 min with the applied waveform repeatedly cycled at a rate of 10Hz. Then a series of stimulations was performed, 2-3 stimulations of each parameter: pulse trains of 30Hz 15 total pulses (p), 60Hz 30p, or 60Hz 60p. This was repeated 2 additional times, for a cumulative dose of 1.5 mg/kg METH or saline given to the rat by the end of the session. Vaginal smears were taken every day and before FSCV testing occurred except during surgery recovery days. When the ES DA response was examined in females who were in different stages of the estrous cycle, the majority of females were in diestrus and no differences between stages were found.

**Figure 2.**
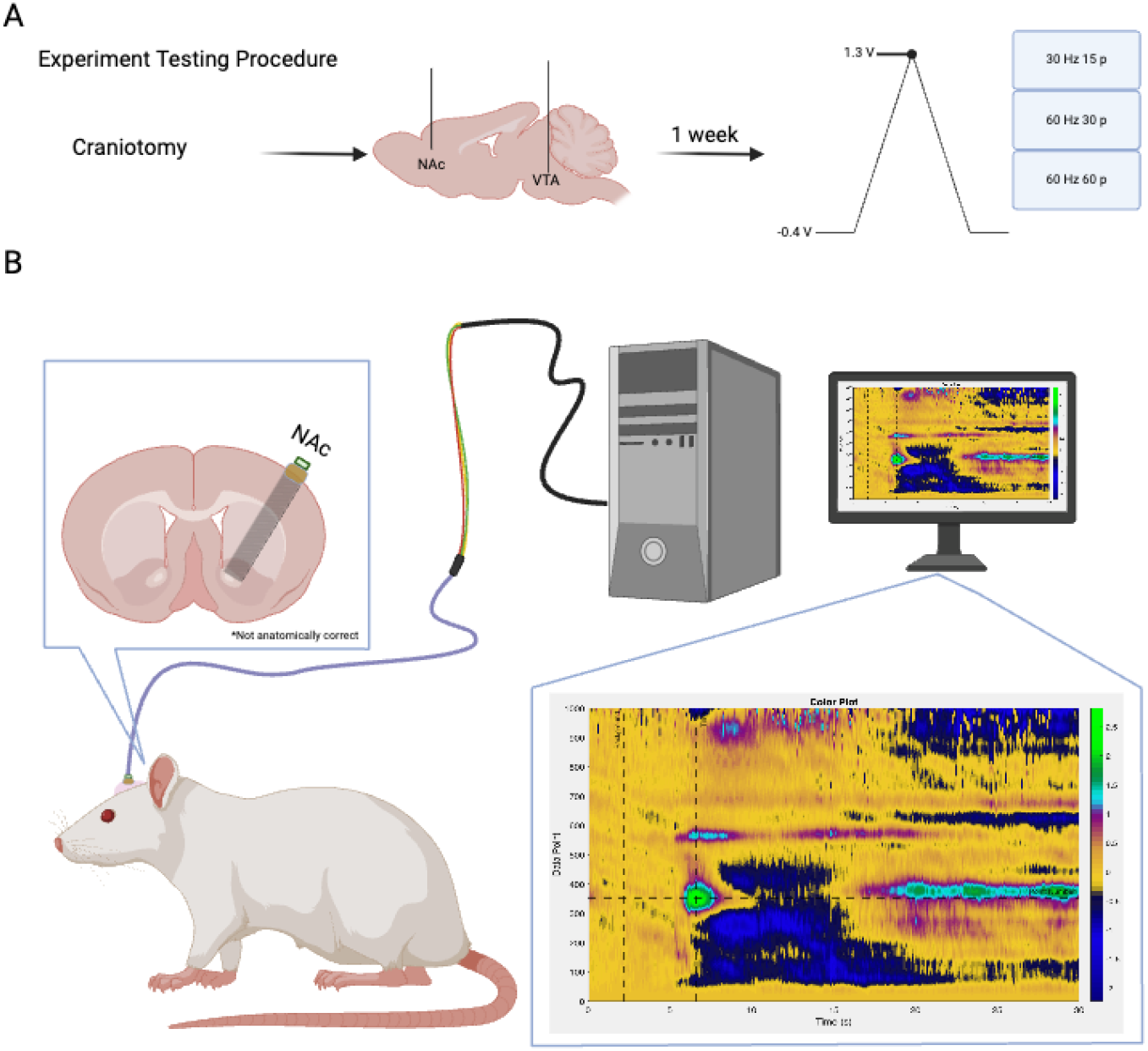
Testing procedure flow diagram. **A**. 16-channel FSCV electrodes were planted into the NAc and stimulating electrodes were implanted into the VTA as described. **B**. Animals were connected to the acquisition system via a headstage and cable during behavioral experiments. Created with BioRender.com

### Data Processing

All FSCV data was collected and stored as described previously [21]. Principle component analysis was performed using an in-house MATLAB code. A standard training set was constructed based on *in vivo* recordings. Current was converted to concentration using the electrode calibration factor obtained before implantation. If the calculated concentration for a channel fell outside of the mean of all 16 channels ± 2 standard deviations, it was identified as an outlier and removed. Custom MATLAB code is available from the corresponding author upon request.

### Statistical Analysis

All statistical analyses were performed using GraphPad Prism v10.0. Data was analyzed using a 2-way ANOVA to look at the effects of maximum increase in DA (mean max DA was defined as the difference between DA prior to stimulation and the peak DA concentration in nM within 10 sec of the stimulation pulse). Group effects in ES DA in the NAcC and NAcS were compared. To look at significant differences of sex and social housing, a Tukey’s comparison test was used for post-hoc tests.

### Histology

Slice-in-place procedures were used to maintain the placement of individual electrode fibers in the brain. The skull was extracted post-mortem with the implant intact and the entire skull was decalcified and then sectioned as previously described [20,21]. Staining protocols were described previously [20,21]. Antibodies: primary antibodies: mouse anti-NeuN (Millipore, MAB277 lot 3574318, 1:500 dilution), chicken anti-tyrosine hydroxylase (TH) (Abcam, AB76422 lot GR3367897, 1:500 dilution), rabbit anti-calbindin (Swant, CB-28a lot 9.20, 1:500 dilution) or rabbit anti-μ-opioid receptor (M-OR) (Immunostar, #24216 lot 20290001, 1:500 dilution); secondary antibodies: donkey anti-mouse Alexa 647 (Jackson, 715-605-150, 1:500 dilution), anti-chicken Alexa 488 (Jackson, 703-545-155, 1:500 dilution), anti-rabbit Alexa 546 (Life Technologies, A10040, 1:500 dilution), DAPI (Invitrogen, D1306, 1:2000 dilution).

### Histology - Imaging and Analysis

Sections were imaged on a Zeiss LSM 780 confocal microscope using a 10x objective (Zeiss, C-Apochromat 10x/0.45 W M27). Sections were cleared by immersion in 80% 2,2’-thiodiethanol (TDE) (Sigma, 88561) for 45 minutes to 1 hour and mounted on electrostatic slides with slide spacers containing TDE. The active surface of the working electrodes is a cylinder with a height of ∼50 μm and a diameter of approximately 6.8 μm. A z-stack image containing the full 16 fibers with 8 μm steps was used to locate each electrode tip. Images were analyzed using the Fiji distribution of ImageJ [22]. Electrodes targeting the NAc were assigned to either the NAcC, NAcS or not in either (excluded) based on staining surrounding the electrode tips as described previously [21]. Briefly, the NAc core was defined by the presence of TH and calbindin staining, the NAcS was defined by the presence of TH staining without calbindin. Electrodes in areas without TH staining were excluded.

## Results

Baseline ES DA release on the first day of testing from these animals was reported previously [21]. To summarize, in the NAcC pair-housed females had higher ES DA release at 60 Hz, 30p compared to the single females, single-housed males and pair-housed males. There were no other differences among the groups. At 60Hz 60p, pair-housed females had significantly more ES DA release compared to the single females. In the NAcS, there were no significant differences among groups found at 60Hz 30p or 60Hz 60p. In pairwise comparisons, there was a significant difference between core and shell in social housed females. No other group differences were found at any stimulation parameter [21]. There was substantial variability among fibers within a brain region and across groups. The coefficient of variation (STDEV/mean) was above 0.5 for all groups and close to 1.0 for many groups. This variability may underlie the failure to find a difference between NAcC vs. NAcS under baseline conditions.

### 60Hz 30p – ES DA Across Weeks: NAcC

A two-way ANOVA was conducted to analyze the effects of social housing on ES DA in the NAcC at 60Hz 30p after each METH injection. At 0.5 mg/kg METH there was an effect of social housing in females ((F (1, 265) = 4.601; p = 0.0329; Fig. 3A)) and an effect of time ((F (2, 265) = 4.562; p = 0.0113; Fig. 3)), such that socially housed females exhibited a greater increase in ES DA release in the NAcC than did single females over time.

**Figure 3:**
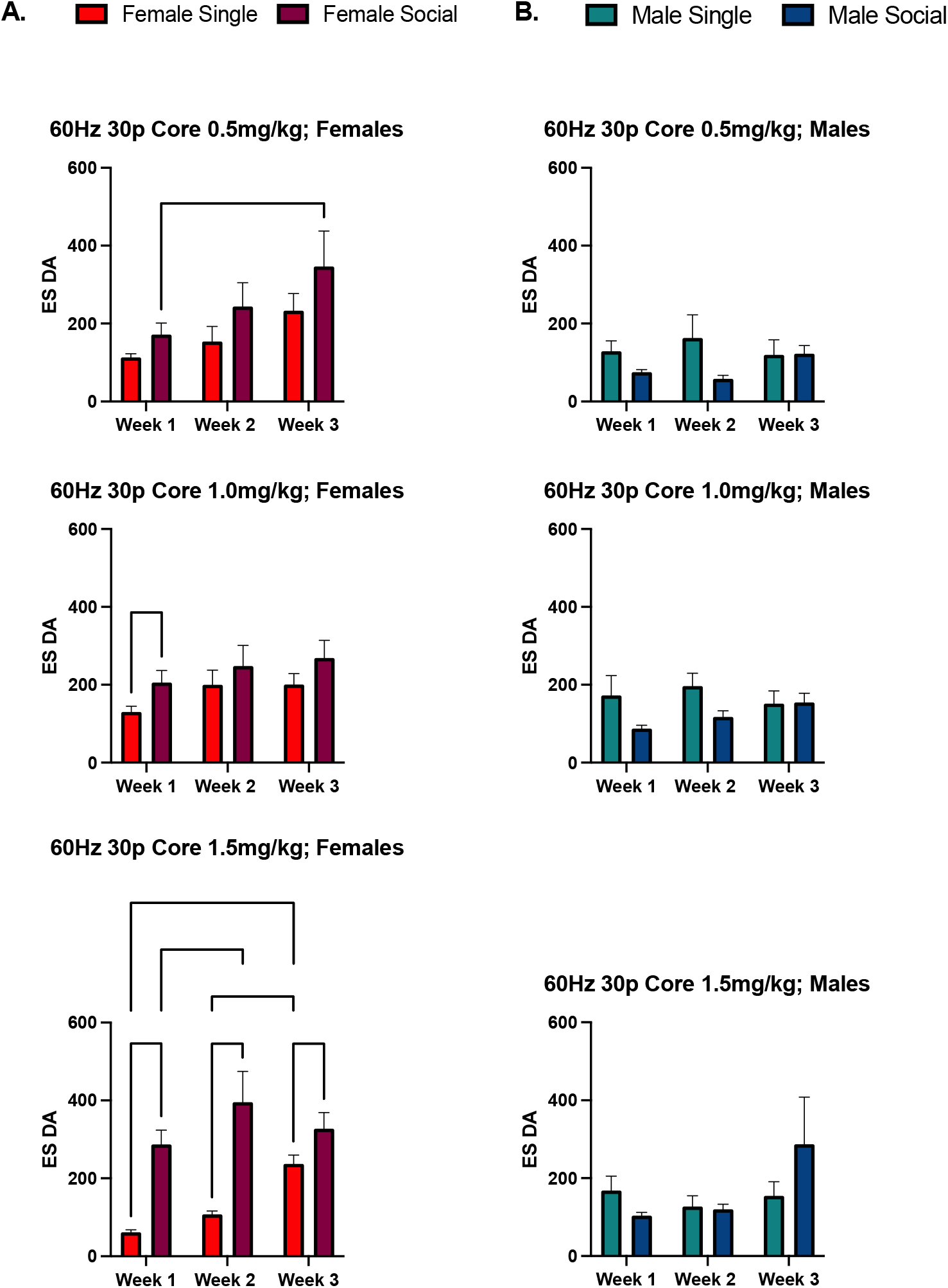
60Hz 30p; NAc Core averages across weeks. **A. Females in the NAcC**, 60Hz 30p: Socially housed females had higher ES DA compared to single housed females. At 1.5 mg/kg METH (Inj 3) single housed females had an increase ES DA across the weeks while socially housed females did not. **B**. Males in the NAcC, 60Hz 30p: No significant differences were found. Bars shown indicate SEM. *p < 0.05, ***p < 0.001.****p < 0.0001.

**Figure 4:**
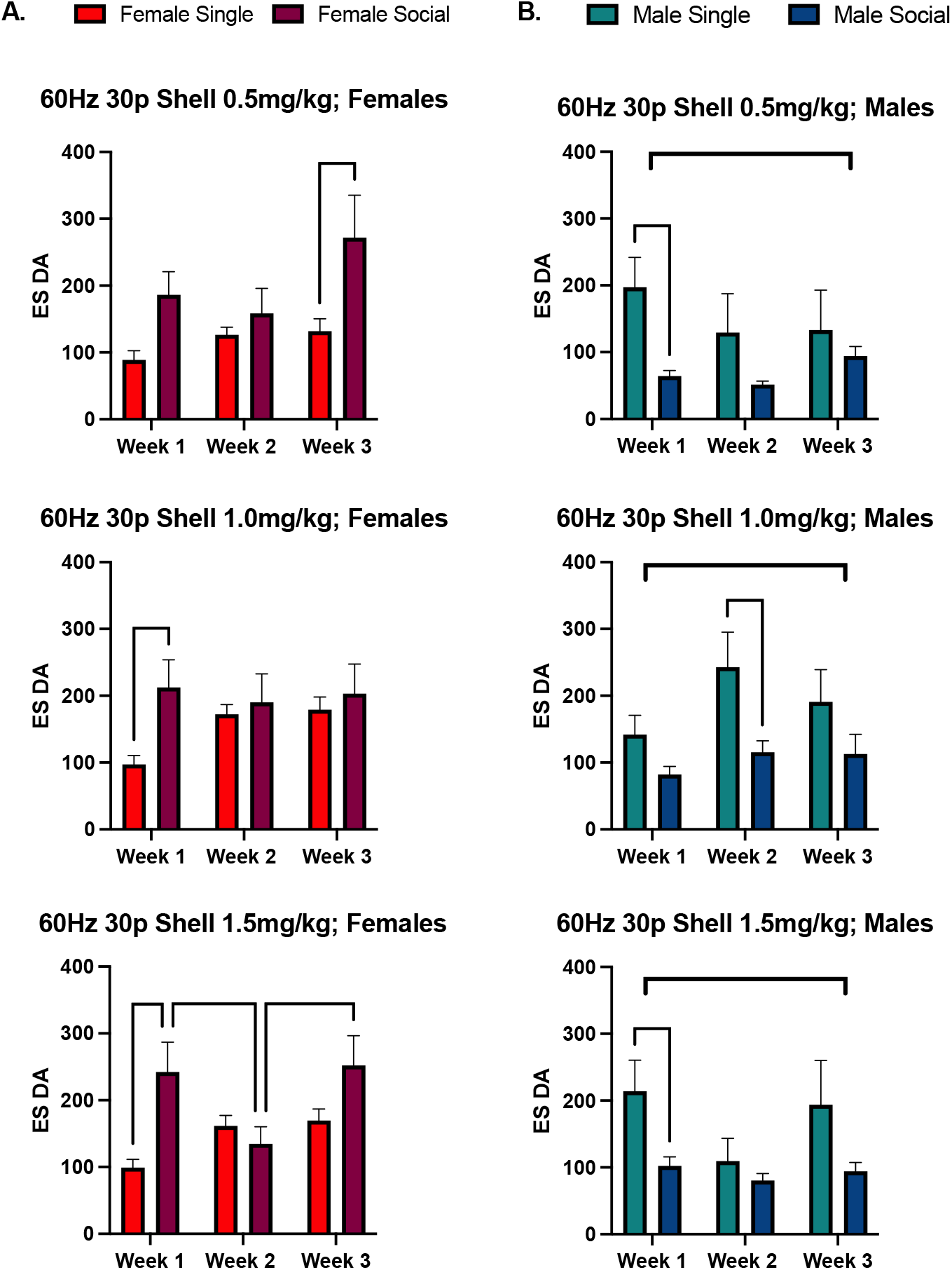
60Hz 30p; NAcS averages across weeks. **A. Females in the NAcS**, 60Hz 30p: Socially housed females had higher ES DA compared to single housed females at 0.5 mg/kg METH and 1.5 mg/kg METH. At 1.5 mg/kg METH, socially housed females had an increase in ES DA across the weeks while single housed females did not. **B. Males in the NAcS**, 60Hz 30p: There was an effect of social housing, where socially housed males had higher ES DA than single housed males. Bars shown indicate SEM. *p < 0.05, **p < 0.01

At 1.0 mg/kg METH in the NAcC socially housed females had greater ES DA release than the single females at week 1 ((F (1, 257) = 5.995; p = 0.0150; Fig. 3A)). At 1.5 mg/kg METH, in females there was also an effect of social housing, with socially housed females exhibiting greater ES DA release in the NAcC than single females ((F (1, 286) = 89.49; p < 0.0001; Fig. 3A)). There was also a significant increase in the average ES DA release in the NAcC across weeks ((F (2, 286) = 11.68; p < 0.0001; Fig. 3A)).

In males, no effect of social housing or time on DA release in the NAcC was found at any of the three METH cumulative doses (Fig. 3B).

#### Dose-response

A two-way ANOVA was used to analyze how ES DA release in the NAcC changed after each METH injection in each week for females and males. In females, at week 1, there was an effect of social housing ((F (1, 298) = 54.21, p < 0.0001)) and an interaction ((F (2, 298) = 6.731, p = 0.0014)). There was an effect of social housing in week 2 ((F (1, 268) = 11.48, p = 0.0008)) and week 3 ((F (1, 242) = 5.550, p = 0.0193)), where socially housed females had the higher ES DA in the NAcC than single females. In males, at week 1there was an effect of social housing ((F (1, 199) = 4.210, p = 0.415)), where socially housed males had higher ES DA in the NAcC than single males. No effects were found at week 2 and week 3 (Figure 3B).

### 60Hz 30p – ES DA Across Weeks: NAcS

A two-way ANOVA was conducted to analyze the effects of social housing on DA release in the NAcS in females and males in ES DA after each METH injection. At 0.5 mg/kg METH there was an effect of social housing in females ((F (1, 234) = 6.584; p = 0.0109; Fig. 3A)) and no change over time. In pairwise comparisons there was greater ES DA from the NAcS of social females vs. single females on week 3. At 1.0 mg/kg METH there was no effect of social housing in females nor an effect of time. Lastly, at 1.5 mg/kg METH there was an effect of social housing ((F (1, 213) = 7.206; p < 0.0078; Fig. 3A)), and a significant difference in the ES DA average across time ((F (1, 236) = 5.117; p = 0.0246; Fig. 3A)). At week 1 and week 3 there was greater ES DA from the NAcS of social females vs. single females. Socially housed males had greater ES DA release in the NAcS than single males after 0.5 mg/kg METH injection F (1, 300) = 5.602, p = 0.0186; Fig. 20B), 1.0 mg.kg METH (1.0mg/kg) ((F (1, 223) = 7.795, p = 0.0057; Fig. 20B), and 1.5 mg/kg METH ((F (1, 333) = 5.688, p = 0.0176; Fig. 3B)), with no effect across time at any dose.

#### Dose-response

In females, in week 1 there was an effect of social housing ((F (1, 219) = 9.571, p = 0.0022)) where socially housed females had higher ES DA release in the NAcS than single housed females, and there was also an interaction ((F (2, 219) = 0.7305, p = 0.4828)). No effects were found at week 2 and week 3. In males there was an effect of social housing in week 1 ((F (1, 330) = 10.84, p = 0.0011)) and week 2 ((F (1, 243) = 6.178, p = 0.0136)), single housed males had higher ES DA release in the NAcS than socially housed males. No effects were found at week 3.

### 60Hz 60p – ES DA Across Weeks: NAcC

A two-way ANOVA was conducted to analyze the effects of social housing in females in ES DA in the NAcC after each METH injection. At 0.5 mg/kg METH there was no effect of social housing or across weeks in females (Figure 5A). At 1.0 mg/kg METH there was an effect of social housing ((F (1, 144) = 19.99; p < 0.0001)), where socially housed females had higher ES DA in the NAcC compared to single housed females in week 1 & week 2. The average ES DA in the NAcC each week increased in single housed females ((F (2, 144) = 6.449; p = 0.0021)), showing sensitization of ES stimulated DA release over time (Fig 5A). Lastly, at 1.5 mg/kg METH in females there was an effect of social housing on ES DA release in the NAcC ((F (1, 122) = 9.259; p = 0.0029)), and a significant difference in the ES DA average release across weeks ((F (2, 122) = 5.181; p = 0.0069)).

**Figure 5:**
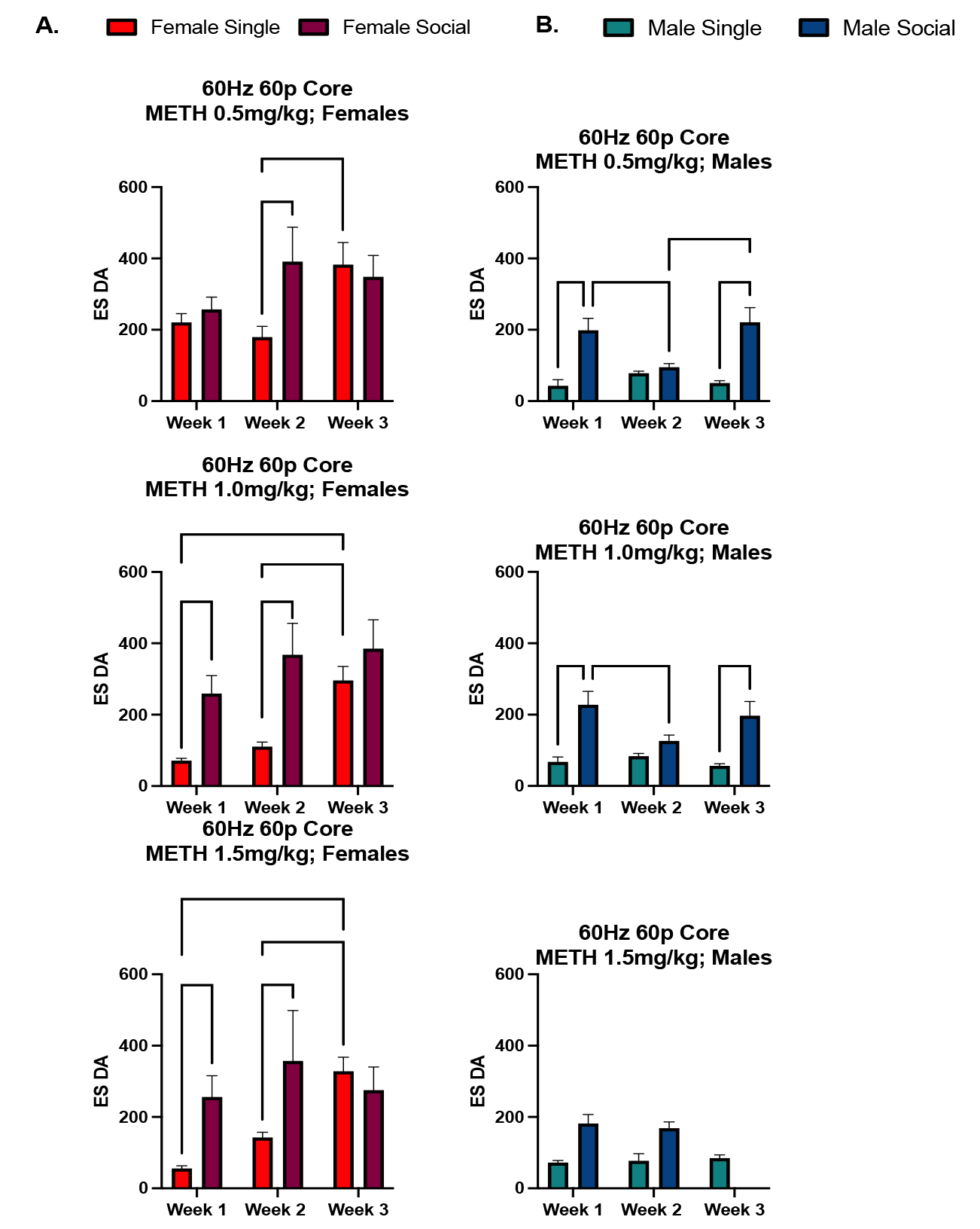
60Hz 60p NAcC averages across weeks. **A. Females in the NAcC**, 60Hz 60p: Socially housed females had higher ES DA compared to single housed females at 1.0 mg/kg METH and 1.5 mg/kg METH. At 1.0 mg/kg METH and 1.5 mg/kg METH single housed females had an increase ES DA across the weeks while socially housed females did not. **B. Males in the NAcC**, 60Hz 60p: There was an effect of social housing, where socially housed males had higher ES DA than single housed males at 0.5 mg/kg METH and 1.0 mg/kg METH.

A two-way ANOVA was then conducted to analyze the effects of social housing in males in ES DA in the NAcC after each METH injection. In males, at 0.5 mg/kg METH there was an effect of social housing on ES DA release in the NAcC where socially housed males had greater ES DA release from the NAcC than single males ((F (1,85) = 2.050, p < 0.0001)). There was no difference across weeks (Figure 5B). In males, at 1.0 mg/kg METH there was no effect across the weeks, but there was an effect of social housing ((F (1, 102) = 33.94, p < 0.0001)). In males at 1.5 mg/kg METH, there was an effect of social housing ((F (1, 101) = 46.66, p < 0.0001)) however, in the socially housed males the data could not be analyzed due to loss of data in week 3, so the weeks could not be compared.

### 60Hz 60p – ES DA Across Weeks: NAcS

A two-way ANOVA was conducted to analyze the effects of social housing in females and males in ES DA in the NAcS after each METH injection. At 0.5 mg/kg METH there was no effect of social housing, but there was a significant difference in the ES DA average across weeks (Figure 6A;(F (2, 162) = 3.309; p < 0.0390)). At 1.0 mg/kg METH there was no effect of social housing but again there was a significant difference in the ES DA in the NAcS average across weeks ((F (2, 249) = 4.978; p < 0.0076)). Lastly, at 1.5 mg/kg METH there was an effect of social housing ((F (1, 213) = 7.206; p < 0.0078)), and a significant difference in the ES DA in the NAcS average across weeks ((F (2, 213) = 4.559; p < 0.0115)).

**Figure 6:**
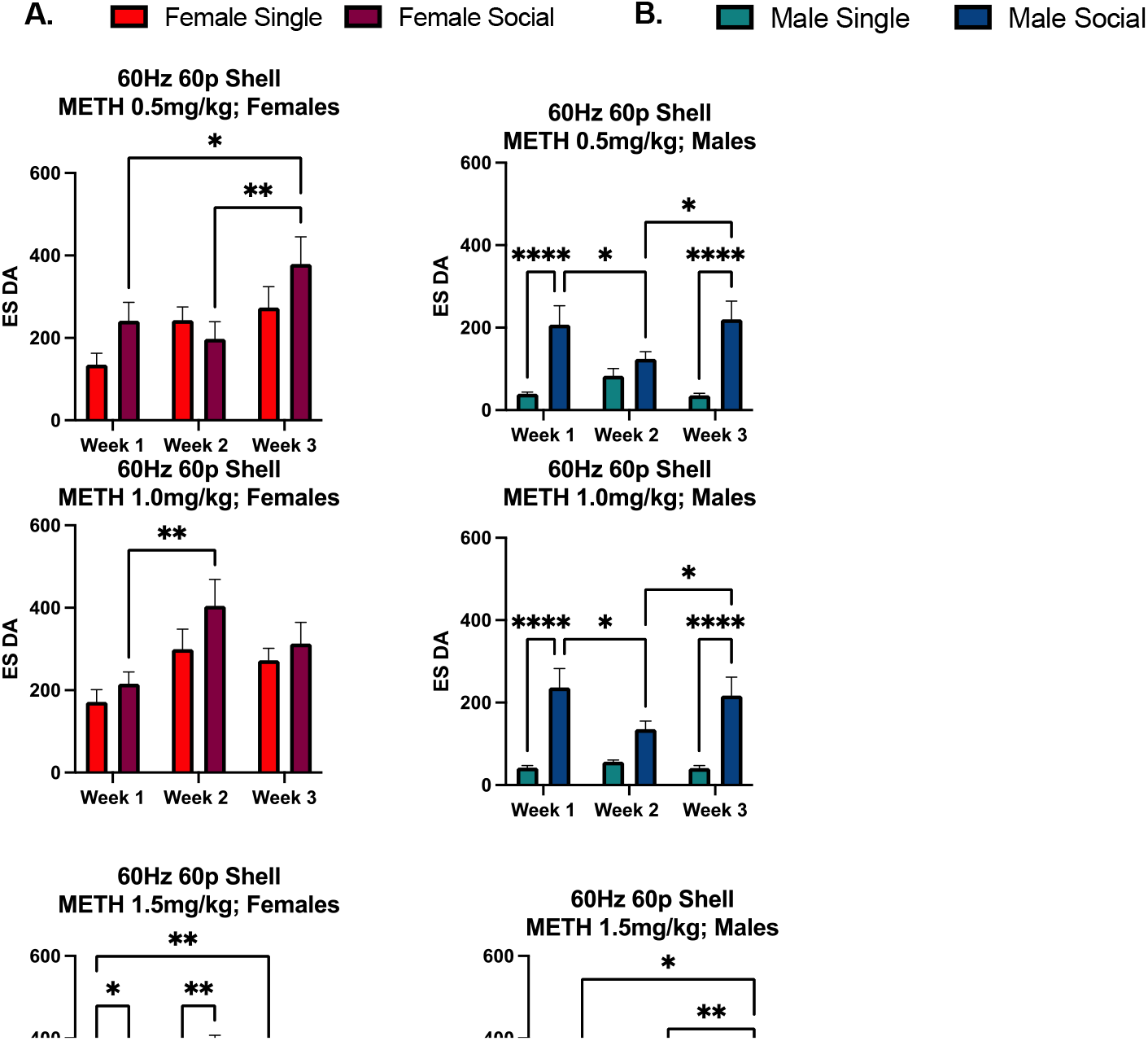
60Hz 60p NAcS averages across weeks. **A. Females ES DA in NAcS**, 60Hz 60p: Socially housed females had higher ES DA compared to single housed females at METH injection 3. At METH injection 3, socially housed females had an increase ES DA across the weeks while single housed females did not. **B. Males ES DA in the NAcS**, 60Hz 60p: There was an effect of social housing, where socially housed males had higher ES DA than single housed males. *p < 0.05, **p < 0.01, ****p < 0.0001.

In males there was an effect of social housing in ES DA release in the NAcS at 0.5mg/kg METH ((F (1, 166) = 28.89, p < 0.001), 1.0 mg/kg METH (1.0mg/kg) ((F (1, 200) = 52.69, p < 0.001), and 1.5 mg/kg METH ((F (1, 174) = 39.47, p < 0.001). There was no effect across weeks at any dose. However, there was an interaction of weeks and social housing at 1.5 mg/kg METH ((F (2, 174) = 4.263, p = 0.0156)), where the socially housed males decreased in ES DA as the single housed males increased in week 3 (Figure 6B).

### Dose-Response Comparisons

#### NAcC

Data were analyzed to determine if there was an effect of the dose of METH on ES DA release from the NAcC in single vs. socially housed rats. At week 1 ((F (1, 298) = 54.21, p < 0.0001)) and week 2 ((F (1, 268) = 11.48, p = 0.0008)) there was an effect of social housing ((F (1, 298) = 54.21, p < 0.0001)), where socially housed females had higher ES DA than single housed females. There no effects found in week 3. In males, at week 1 ((F (1, 90) = 37.25, p < 0.0001)) and week 2 ((F (1, 85) = 20.55, p < 0.0001)), there was an effect of social housing.

Socially housed males had higher ES DA in the NAcC compared to single housed males. At week 2, there was also a dose effect ((F (2, 85) = 3.580, p = 0.0322)), ES DA in the NAcC increased after each METH injection and there was an interaction ((F (2, 85) = 3.967, p = 0.0225)) of social housing and dose at 0.5 mg/kg METH. Week 3 could not be analyzed as 1.5 mg/kg METH is missing for socially housed males.

### NAcS

Data were analyzed to determine if there was an effect of the dose of METH on ES DA release from the NAcS. In females, at week 1 there were effects of social housing F (12, 219) = 9.571, p = 0.0022 where socially housed females had higher ES DA compared to single housed females. There were no effects found in week 2 & 3. In males there was an effect of social housing at week 1 ((F (1, 190) = 58.30, p < 0.0001)), week 2 ((F (1, 154) = 37.94, p < 0.0001)), and week 3 ((F (1, 196) = 32.76, p < 0.0001)), where socially housed males had higher ES DA compared to single housed males. At week 2, there was an interaction ((F (2, 154) = 6.458, p = 0.0020)) at 0.5 mg/kg METH and at week 3 there was an interaction ((F (2, 196) = 3.556, p = 0.0304)) at 1.5 mg/kg METH.

### Regional Differences

#### 60Hz 60p

Differences between NAcC and NAcS were compared at 60Hz60p for single females (Figure 7). In single females, at 0.5 mg/kg METH at 60Hz60p there was an effect of the average ES DA across weeks ((F (2, 118) = 6.169, p = 0.0028)) but no differences in the brain regions and an interaction ((F (2, 118) = 3.273, p = 0.0414. At 1.0 mg/kg METH there was effect of the average ES DA across weeks ((F (2, 179) = 16.07, p < 0.0001)), in both brain regions ((F (1, 179) = 13.11, p = 0.0004)), and an interaction ((F (2, 179) = 6.294, p = 0.0023)). At 1.5 mg/kg METH there was an effect of the average ES DA across weeks ((F (2, 286) = 41.65, p < 0.0001) and there was a significant difference in the brain regions, ((F (1, 286) = 18.06, p < 0.0001)). The single housed females showed increased sensitization over time, making them more vulnerable to addictive-like behaviors. No effects were found in socially housed females.

**Figure 1:**
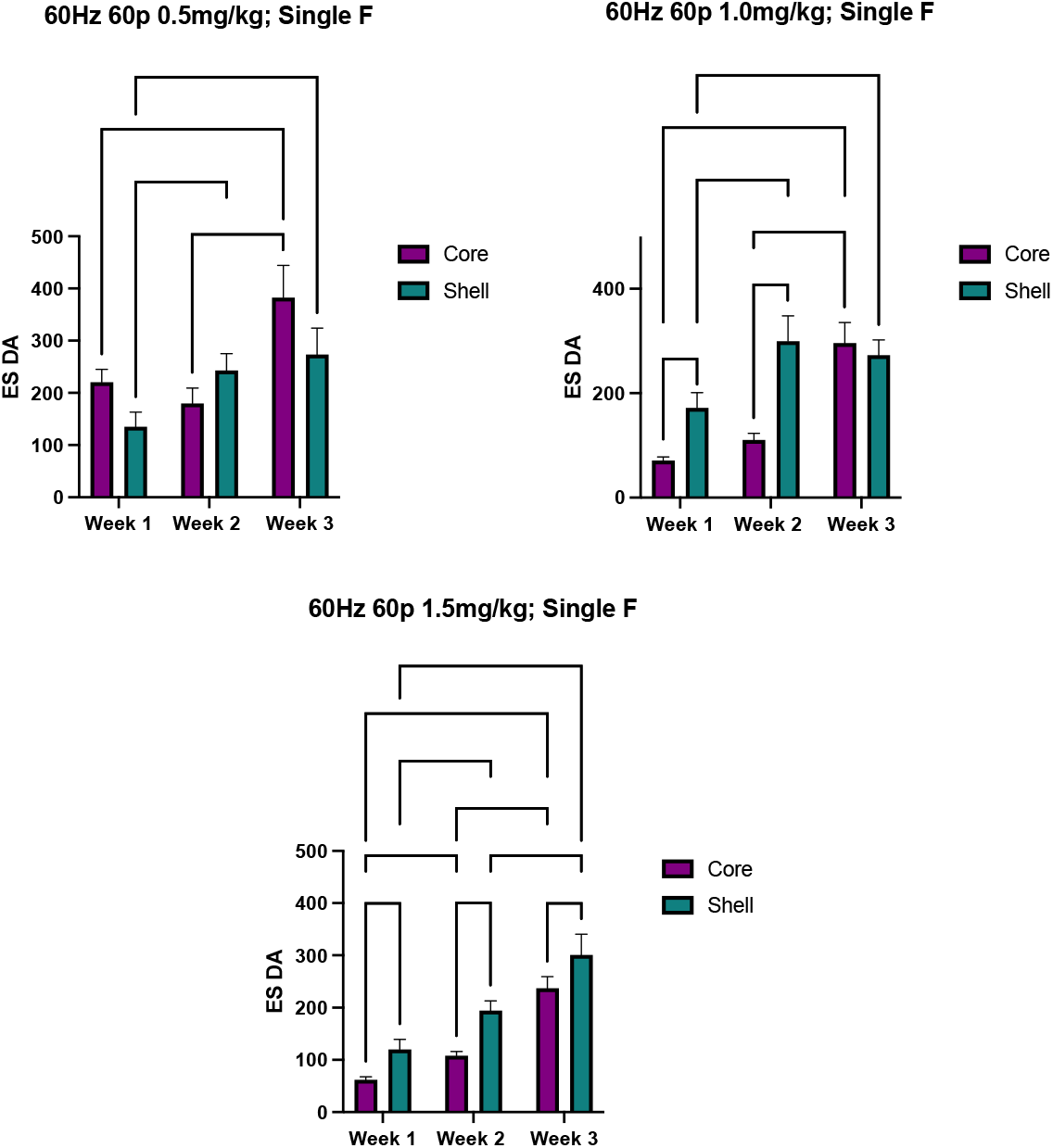
Regional ES DA differences in single housed females.

There was an effect found in females across all three METH injections found where the NAcS ES DA was higher than the NAcC ES DA. *p < 0.05, **p < 0.01, *** p < 0.001, ****p < 0.0001.

Differences between NAcC and NAcS were compared at 60Hz60p for single males (Figure 8). In single housed males, there was an effect of weeks at 0.5 mg/kg METH ((F (2, 133) = 14.51, p < 0.0001)), 1.0 mg/kg METH ((F (2, 184) = 6.528, p = 0.0018)) and 1.5 mg/kg METH ((F (2, 179) = 4.037, p = 0.0193)). There was significant differences between the brain regions at 1.0 mg/kg METH ((F (1, 184) = 16.94, p < 0.0001)) and 1.5 mg/kg METH ((F (1, 179) = 53.28, p < 0.0001)). No effects were found in socially housed males.

**Figure 2:**
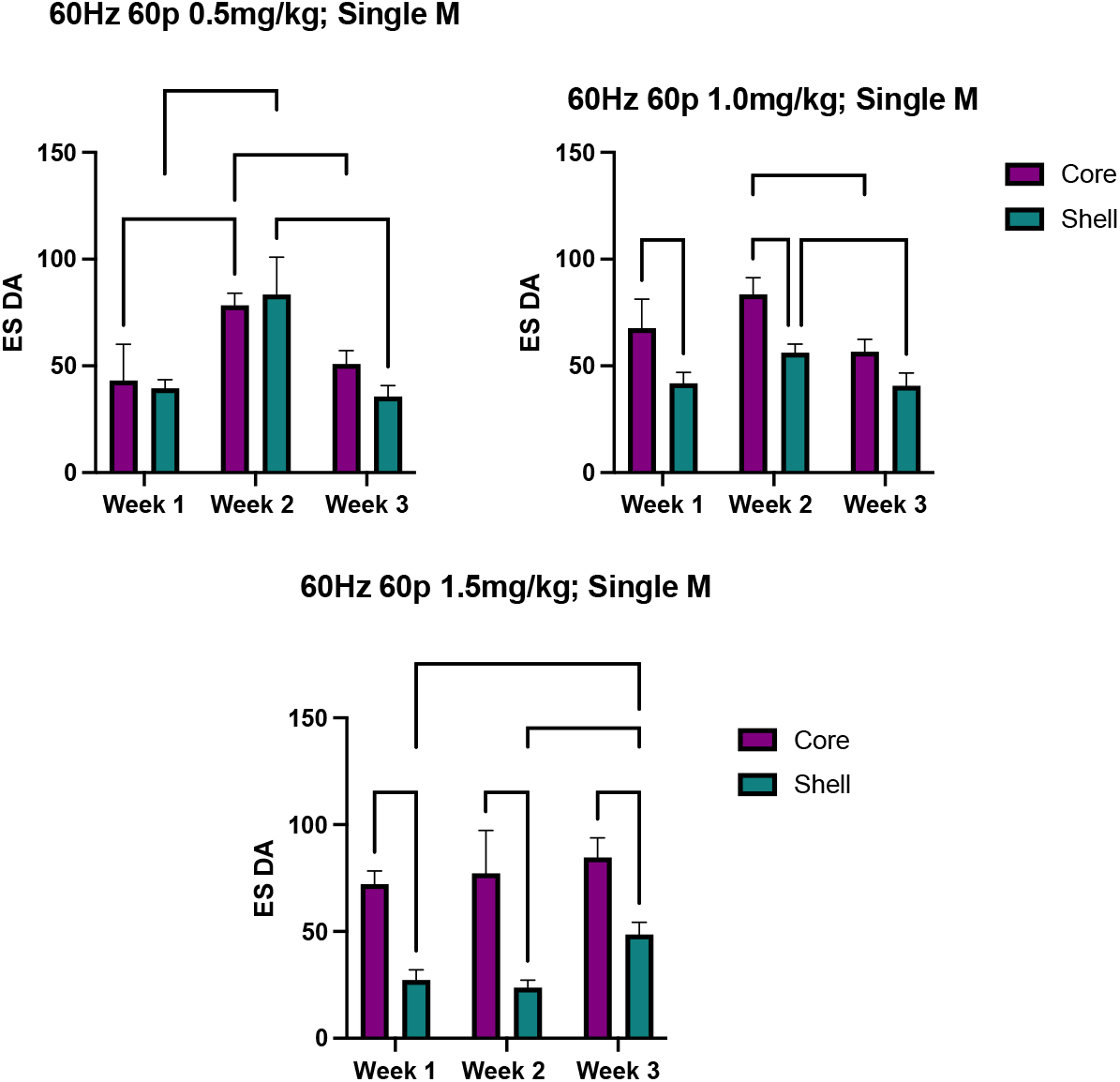
Region differences in single housed males. There was an effect found in males across at METH injection 2 and METH injection 3 where the NAcC ES DA was higher than the NAcS ES DA. *p < 0.05, **p < 0.01, *** p < 0.001, ****p < 0.0001.

**Figure 9:**
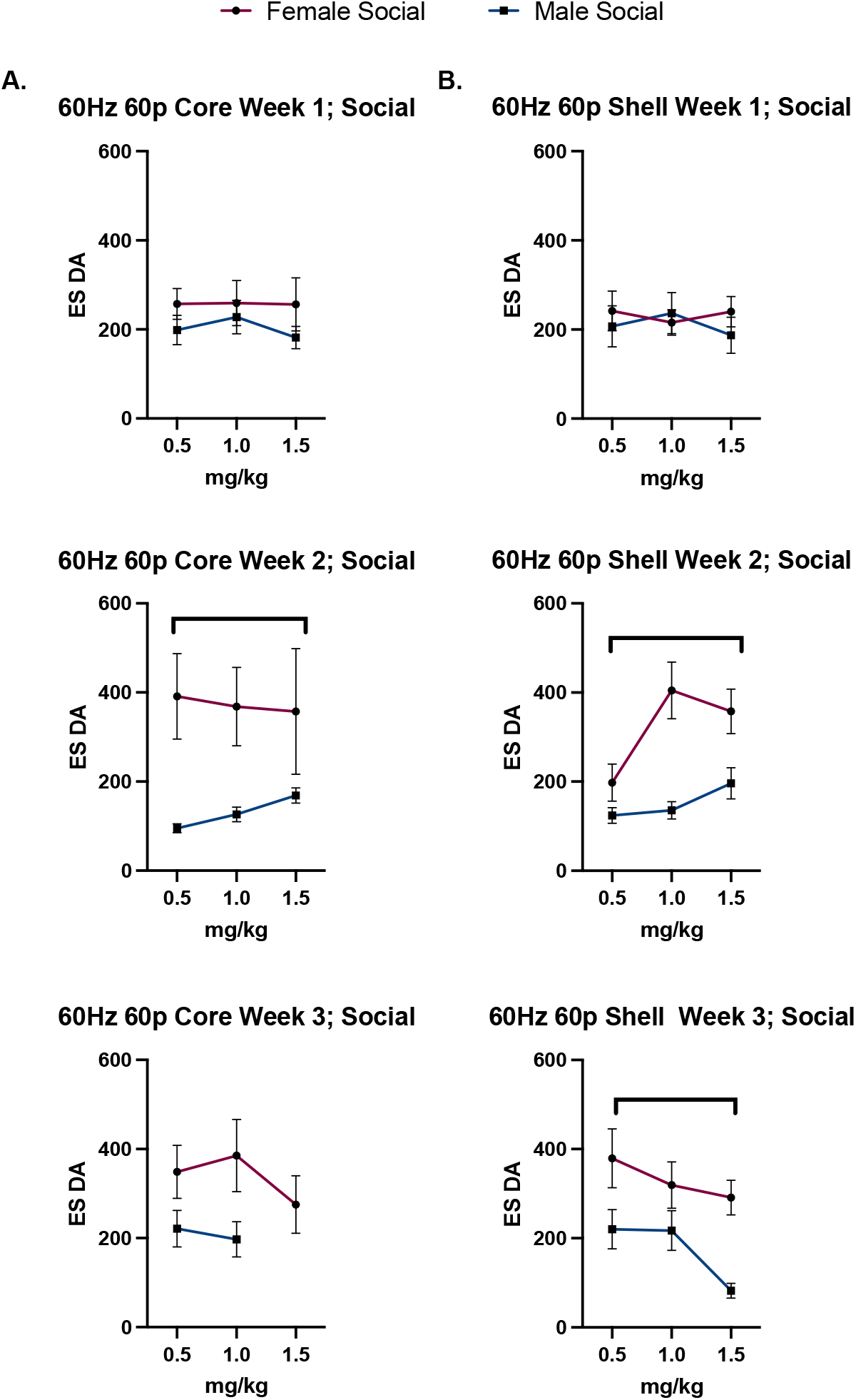
Sex differences in the socially housed animals. **A**, NAcC, 60Hz 60p: Socially housed females had higher ES DA compared to socially housed females at week 2. No dose effects found. **B**, NAcS, 60Hz 60p: Socially housed females had higher ES DA compared to socially housed males at week 2 and week 3. No dose effects found. Bars shown indicate SEM. **p < 0.01, *** p < 0.001,

**Figure 10:**
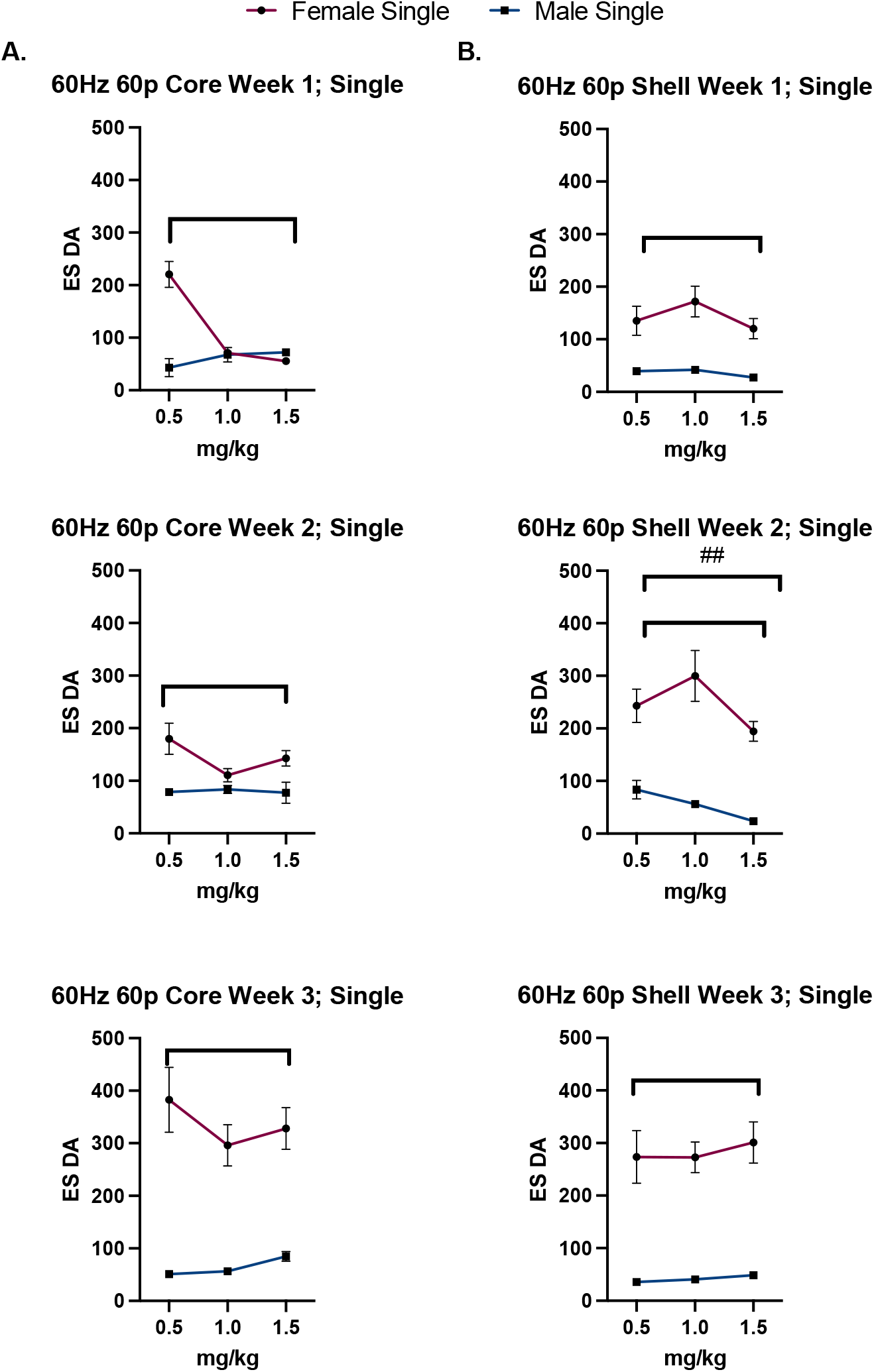
Sex differences in the single housed animals. **A**, NAcC, 60Hz 60p: Single housed females had higher ES DA compared to single housed males across all weeks. No dose effects found. **B**, NAcS, 60Hz 60p: Single housed females had higher ES DA compared to single housed males across all weeks. There was a dose effects found in week 2, where ES DA decreased after each METH injection. Bars shown indicate SEM. **p < 0.01, *** p < 0.001, ****p < 0.0001, ## p < 0.01 (dose-effect).

### Sex Differences

#### Socially housed rats

In week 1, there was no difference between socially housed females and males in for the NAcC or the NAcS (Fig. 25; A & B). In week 2, socially housed females had higher ES DA in the NAcC compared to socially housed males ((F (1, 80) = 9.526, p = 0.0028; Fig. 25A)). In week 2, socially housed females had higher ES DA in the NASh compared to socially housed males ((F (1, 205) = 13.48, p = 0.0003; Fig. 25B)). Lastly, in week 3 there were no significant differences found in the NAcC. In week 3 in the NAcS, socially housed females had higher ES DA compared to socially housed males ((F (1, 188) = 9.863, p = 0.0020; Fig. 25B)).

#### Single housed rats

In week 1, single housed females had higher ES DA in the NAcC ((F (1, 110) = 29.52, p < 0.0001; Fig. 26A)) and the NAcS ((F (1, 188) = 60.53, p < 0.0001; Fig. 26B)) compared to single housed males. There was also an interaction where single housed males had higher ES DA at the 1.5mg/kg injection in the NAcC ((F (2, 110) = 27.54, p < 0.0001; Fig. 26A)). Lastly, there was a decrease in ES DA in both females and males across each injection ((F (2, 110) = 13.85, p < 0.0001; Fig. 26A)). In week 2, single housed females had higher ES DA in the NAcC ((F (1, 147) = 121.1, p < 0.0001; Fig. 26A)) and the NAcS ((F (1, 153) = 95.92, p < 0.0001; Fig. 26B)) compared to single housed males. Also in the NAcS, there was a decrease in females and males from 0.5mg/kg injection to the last injection, 1.5m/kg ((F (2, 153) = 5.830, p = 0.0036; Fig. 26B)). Lastly, in week 3 single housed females had higher ES DA in the NAcC ((F (1, 151) = 11.71, p < 0.0001; Fig. 26A)) and the NAcS ((F (1, 208) = 161.1, p < 0.0001; Fig. 26B)) compared to single housed males.

## Discussion

The results showed that social housing enhances electrically stimulated DA release in males and that there was greater DA release in NAc core than shell in single males, but no difference in socially housed males. In females, social housing also enhanced ES DA release. In single females there was greater ES DA release in shell than in core.

In single housed females there was greater ES DA release over time, while the socially housed females and males had high ES DA release that remained stable over time. These results suggest that social housing protects females from sensitization, making single females more vulnerable to the addictive properties of METH. Other studies using cocaine IntA self-administration have shown that females sensitize but not males [23].

Notably, there were social housing differences in both females and males. In females, socially housed females had consistently higher ES DA release compared to single housed females. Before METH is introduced socially housed females have higher ES DA compared to single housed females. After METH, socially housed females, the ES DA remained high for majority of the METH injections and across time, suggesting that there is a ceiling effect which protects socially housed females from the addictive-like properties of METH. This result did not support the initial hypothesis that single housed females would have higher ES DA compared to socially housed females. Males did have a social housing differences in the NAcC and the NAcS at the 60Hz 60p parameter but there were not consistent differences with other stimulation parameters.

There are some limitations to this study, fibers did not consistently record DA or broke over time leading to missing data points. This did not allow us to compare individually fibers over time so further studies could focus on focusing on analyzing fibers over time. Another limitation is that the animals were given an injection through an IP injection, making it difficult to determine exactly how much METH was absorbed into the body. Future studies should consider using an intravenous catheter to administer METH to be more precise in the overall METH exposure.

## Conclusion

The present study showed that socially housed females had higher ES DA than single housed females, but this increase in higher ES DA may be a protective factor for females. Single housed females had sensitization across time, making them more vulnerable to addictive-like behaviors. While socially housed females could potentially have a ceiling affect, protecting them from these sensitization effects. Single males and social housed males did not have the same sensitization effect.

## Acknowledgement

The research reported here was supported by a grant from the USPHS to JBB. 1R01DA046403 Becker (PI) NIH/National Institute on Drug Abuse: Social support, oxytocin and motivation for methamphetamine

